# PRY-1/Axin signaling regulates lipid metabolism in *Caenorhabditis elegans*

**DOI:** 10.1101/289462

**Authors:** Ayush Ranawade, Avijit Mallick, Bhagwati P Gupta

## Abstract

The nematode *Caenorhabditis elegans* constitutes a leading animal model to study how signaling pathway components function in conserved biological processes. Here, we describe the role of an Axin family member, *pry-1*, in lipid metabolism. As a central component of the canonical WNT signaling pathway, *pry-1* acts as a scaffold for a multiprotein destruction complex that negatively regulates the expression of WNT target genes. Genome-wide transcriptome profiling of a *pry-1* mutant revealed genes associated with aging and lipid metabolism such as vitellogenins (yolk lipoproteins), fatty acid desaturases, lipases, and fatty acid transporters. Consistent with these categorizations, we found that *pry-1* is crucial for the maintenance of lipid levels. Knock-downs of *vit* genes in a *pry-1* mutant background restored lipid levels, suggesting that vitellogenins contribute to PRY-1 function in lipid metabolic processes. Additionally, lowered expression of desaturases and lipidomics analysis provided evidence that fatty acid synthesis is reduced in *pry-1* mutants. Accordingly, an exogenous supply of oleic acid restored depleted lipids in somatic tissues of worms. Overall, our findings demonstrate that PRY-1/Axin signaling is essential for lipid metabolism and involves the regulation of yolk proteins.

## INTRODUCTION

Axin was identified initially as a negative regulator of the WNT-signaling pathway (1). Subsequently, Axin was shown to also be essential in diverse developmental events including embryogenesis, neuronal differentiation, and tissue homeostasis (2), with Axin homologs exhibiting functional conservation throughout metazoans (3). As a scaffolding protein Axin plays a key role in the regulation of canonical WNT pathway function. It contains multiple domains that facilitate homodimerization and interactions with the destruction-complex proteins dishevelled, APC, and GSK-3β (4, 5). In turn, the destruction complex initiates the phosphorylation and consequent proteolysis of the transcriptional regulator β-catenin, which promotes expression of WNT target genes (4, 5). In an ON state, the WNT-initiated signal inhibits the action of the destruction complex, thereby causing cytoplasmic accumulation of β-catenin, which subsequently translocates to the nucleus and promotes the transcription of target genes (4, 6). Constitutive activation of β-catenin, consequent to the loss of destruction-complex function, is often associated with cancers and various other disorders affecting the lungs, heart, muscles, and bones (4). Thus, Axin function appears crucial toward ensuring precise regulation of β-catenin-mediated WNT signaling.

Similar to the mammalian systems, Axin homologs in invertebrates also play crucial role in the regulation of WNT signaling (7). In the nematode *Caenorhabditis elegans*, PRY-1/Axin interacts with APR-1/APC and GSK-3/GSK-3β to regulate BAR-1/β-Catenin-mediated gene transcription (3). *pry-1* mutants show defects that are consistent with overactivation of WNT signaling, such as neuronal differentiation and formation of multiple tissues (7).

Besides its role in development and diseases, WNT pathway components are also involved in cellular senescence, tissue aging, and nutrient metabolic processes. However, the mechanism by which the pathway affects these various processes is not well understood. Here, we investigated the role of *pry-1*/*Axin* in *C. elegans* and provide evidence for its important role in the maintenance of lipid metabolism through the regulation of vitellogenesis.

## EXPERIMENTAL PROCEDURES

### Strains

Worms were grown on standard Nematode Growth Medium (NGM) plates using procedure described previously (8). Cultures were maintained at 20 °C unless mentioned otherwise. The *mu38* allele of *pry-1* was obtained from CGC. All strains were outcrossed at least three times before doing the experiments.

The genotypes of the strains used in this study are: N2 (wild-type), DY220 *pry-1(mu38)I*, PS4943 *huIs[dpy-20;hsp16-2*::*dNT-bar-1]; syIs148*, RB1982 *vit-1(ok2616)X*, RB2365 *vit-2(ok3211)X*, RB1815 *vit-3(ok2348)X*, RB2202 *vit-4(ok2982)X*, RB2382 *vit-5(ok3239)X*.

### Molecular Biology

For qRT-PCR experiments mRNA was extracted from bleach synchronized worms by Tri-reagent (Catalog Number T9424, Sigma-Aldrich Canada) according to the manufacturer’s instructions. cDNA was synthesized from total RNA using oligo (dT) primers and other reagents in the ProtoScript^®^ First Strand cDNA Synthesis Kit (Catalog Number E6300S, NEB, Canada). Quantitative real-time PCR (qRT-PCR) analysis was performed on a CFX 96 BioRad cycler in triplicate with SensiFAST™ SYBR^®^ Green Kit (Catalog Number BIO-98005, USA), according to the manufacturer’s instructions. *pmp-3* was used as a reference gene in all assays. CFX manager was used for the Ct and *p*-value calculations. At least three replicates were performed for each assay.

### RNAi

For RNAi experiments, *Escherichia coli* HT115 expressing target specific dsRNA were grown on plates containing β-lactose (9). Worms were bleach synchronized and seeded onto plates. After becoming young adults, worms were transferred to fresh plates every other day and the numbers of dead worms recorded. For adult specific RNAi, synchronized worms were cultivated on NGM/OP50 plates until the young adult stage and then transferred to the RNAi plates.

### Oil Red O staining

Oil Red O staining was performed as previously reported (10). Briefly, worms were collected from NGM plates, washed with 1× phosphate-buffered saline (PBS) buffer, and re-suspended in 60 µl of 1× PBS (pH 7.4), 120 µl of 2× MRWB buffer (160 mM KCl, 40 mM NaCl, 14 mM Na2-EGTA, 1 mM spermidine-HCl, 0.4 mM spermine, 30 mM Na-PIPES [Na-piperazine-*N, N*′-bis (2-ethanesulfonic acid); pH 7.4], 0.2% β-mercaptoethanol), and 60 µl of 4% paraformaldehyde. The worms were then freeze-thawed 3 times and washed twice with 1× PBS. They were then incubated at room temperature in 60% isopropyl alcohol for 10 minutes for dehydration and stained with freshly prepared Oil Red O solution for at least 48 hours on a shaker. For direct and consistent comparison, all Oil Red O images from the same experiment were acquired under identical settings and exposure times. Animals were mounted and imaged using Q-imaging software and a Micropublisher 3.3 RTV color camera outfitted with DIC optics on a Nikon 80i microscope. NIH ImageJ software was used to quantify Oil Red O intensities (10). A total of 15 to 30 worms were randomly selected from each category in at least two separate batches.

### Brood Assay

Worms were bleach synchronized and allowed to grow to L4 stage for determining the progression of egg-laying and the brood size. Individual worms were picked onto a separate NGM plate with *E. coli* OP50 bacteria and allowed to grow for several days. Worms were repeatedly transferred to a freshly seeded NGM plate and progeny were counted every 24 hours. Data from escaping or dying mothers were omitted from the analyses (11).

### OA supplementation assay

To make OA supplemented NGM agar plates, a 0.1 M water-based stock solution of OA sodium salts (NuCheck Prep, USA) was prepared and stored at –20 °C in the dark. The OA solution was added continuously to the NGM and promptly poured into the plates. The plates were covered with aluminum foil and kept at room temperature overnight to dry. The *E. coli* OP50 strain was seeded onto each plate and allowed to further dry for one to two days in the dark. Oil Red O staining was performed as described above (12).

### Lipase assay

Lipase activity was estimated using commercially available QuantiChrom™ Lipase Assay Kit (BioAssay Systems, USA, Catalog number DLPS-100) and processed according to the manufacturer’s instructions. 1 unit of Lipase catalyzes the cleavage of 1 µmol substrate per minute. Three independent samples of one-day-adult worms were prepared by homogenizing in a 20% glycerol, 0.1 M KCl, 20 mM HEPES (pH 7.6) buffer for further measurements as described earlier (13).

### L1 survival assay

Worms were bleach synchronized and kept in 1.5 ml centrifuge tube. Worms were seeded onto NGM plates approximately 24 hours afterwards regularly for 12 days, and numbers of seeded worms counted. Worms were grown to young adult stage before survivors were counted. The L1 diapause data were statistically compared using an analysis of covariance (ANCOVA) model.

### RNA-Seq and data analysis

*pry-1* targets were examined in synchronized L1 stage animals. At this stage Wnt ligands, receptors, and targets are highly expressed as revealed by microarray studies from SPELL database (14, 15) (Fig. S1A-D). Also, our qRT-PCR experiments showed significant upregulation of three of the Wnt targets, *lin-39*, *egl-5* and *mab-5*, in *pry-1* animals at L1 stage (Fig. S1D). The *pry-1* transcriptome profile can be found in the GEO archive with accession number <u>GSE94412</u>. For RNA-Seq experiments synchronized L1 stage animals were obtained by two successive bleach treatments and RNA was isolated using Trizol-reagent (Sigma, USA, Catalog Number T9424) (16). The quality of total purified RNA was confirmed using bioanalyzer (Agilent 2100 and Nanodrop 1000). cDNA libraries were constructed from 100 - 200ng mRNA using an Illumina-specific commercial kit (TruSeq RNA Sample Preparation Kit v2, Set A Catalog number: RS-122-2001). RNA sequencing was carried out using Illumina Hi-Seq 2000 system at the McGill University Genome Quebec sequencing facility. For each of the N2 and *pry-1(mu38)* strains two biological replicates were used. For each cDNA library, 100 bp paired-end reads were obtained. In total, 30 to 38 million reads were obtained for each sample analyzed for differential gene expression.

The adapters were trimmed using cutadapt/trimgalore, reads with QC values (Phred score) lower than 30 bases were discarded after trimming process (17). Later, processed sequencing reads were mapped to the reference genome (ce6) (UCSC 2013) using the software package Bowtie 1.0.0 (18). 92-95% of total sequenced fragments could be mapped to the genome (Table S2). Transcript-level abundance estimation was performed using eXpress 1.5 software package (19). Among all genes analyzed, 18867 were mapped to known transcripts by at least one sequencing fragment in *C. elegans*. To avoid biases between samples, the gene counts were quantile normalized (17, 20). Using a negative binomial distribution model of DESeq package in R, differentially-expressed genes were called at a false discovery rate (FDR) of 0.05% (21).

GO analysis was carried out with default setting using GoAmigo (*http://amigo.geneontology.org*). A GO-term containing at least three genes with a *p*-value adjusted for multiple comparisons and < 0.05 (Benjamini-Hochberg method) was counted significant (22). Tissue enrichment analysis was performed using Wormbase online TEA tool that employs a tissue ontology previously developed by WormBase (23).

### Gas Chromatography Mass Spectrophotometry (GC-MS)

Fatty Acid analysis protocol was modified from a previously published method (24, 25). Eppendorf tubes, glass vials or any containers used for the extraction process were sonicated in dichloromethane (Catalog number 3601-2-40, Caledon Laboratories Ltd., Canada) for 30 minutes to eliminate lipid contamination. To determine FA composition, few thousand adult worms were collected from three 6-cm plates and washed with sterile water to remove any bacterial contamination. Then the worms were transferred into a screw-capped centrifuge tube and spun at 2,500 RPM for 45 seconds, as much water as possible was removed with a Pasteur pipette and transferred to a Standard Opening (8 mm) glass screw top vials (Agilent, part number 5182-0714) and accurately weigh 50-100 mg of sample. 1 ml of 2.5% H2SO4 (Catalog number 8825-1-05, Caledon Laboratories Ltd., Canada) in methanol (Catalog number 6701-7-40, Caledon Laboratories Ltd., Canada) was added to extract fatty acids from tissues and transmethylate them. Samples were spiked with 10 µl of a recovery standard (stearic acid 120 ng/µl, Catalog number S4751-1G Sigma-Aldrich Canada) and incubated at 80°C in a water bath for an hour. To this, a mixture of 0.2 ml of hexane (Catalog number 3201-7-40, Caledon Laboratories Ltd., Canada) and 1.5 ml of H2O was added and slowly spun to extract fatty acid methyl esters into the hexane layer. Agilent 6890 series gas chromatographer equipped with a 30 × 0.25 mm SP-2380 fused silica capillary column (Supelco USA), helium as the carrier gas at 1.4 ml/minute, and a flame ionization detector was used for FA analysis. Automatic injections of 1 µl samples in the organic phase were made, without splitting, into the GC inlet set to 250 °C. The thermal program began at 50 °C for 2 minutes, then increased linearly to 300° C at a ramping rate of 8 °C/minute and held this temperature for 15 minutes. A constant flow rate of 1 ml/minute helium carrier gas under electronic pressure control was maintained for the fatty acid composition determination by TIC method using standard software. For quantitation of fatty acids, the peaks across all GC-MS runs were aligned using both chromatographic information (retention times) and mass-spectral data (m/z components) to establish the chemical identity of peaks being compared. A relative FA amounts were calculated by dividing each peak area by the sum of areas for all FA peaks appearing on the same chromatogram. For each FA, the quantities determined by GC-MS were successively normalized in two ways: (1) to an internal standard naphthalene-d8, 10 ng/µl (1ng/µl in injection sample) added to each sample prior to sonication and lipid extraction), and (2) to the weight of the samples.

### Statistical Analysis

The statistics were performed using Microsoft Office Excel 2016. If not specifically mentioned, *p* values for the fertility, motility, fat content, L1 survival, and enzyme activity assays were calculated by the Student’s t test after testing for equal distribution of the data and equal variances within the data set. Experiments were performed in triplicate except where stated otherwise. Differences were considered statistically significant at *p* < 0.05, thereby indicating a probability of error lower than 5%. Hypergeometric probability tests and statistical significance of the overlap between two gene sets were done using an online program (*http://nemates.org/MA/progs/overlap_stats.html*).

## RESULTS

### Identification of PRY-1 targets

To gain insights into the mechanism of *pry-1*-mediated signaling, a genome-wide transcriptome analysis was carried out to identify the potential downstream targets. Using RNA-Seq, we identified a total of 2,665 genes (767 upregulated and 1898 downregulated, False Discovery Rate (FDR) *p*-adj 0.05) that were differentially expressed in *pry-1(mu38)* animals during the L1 larval stage (Fig. 1A, Table S3, also see Methods). Of these, the transcription of 1,331 genes was altered twofold or more (FDR, *p*-adj 0.05) (248 upregulated and 1083 downregulated) (Table S3). The average and median fold changes in the expression were 2.2 and 2.0, respectively. Figure 1A shows a scatter plot of all expressed genes. A total of 20 genes was also tested by quantitative polymerase chain reaction (qPCR), which revealed an 85% validation rate (Fig. 1F, G).

**Figure 1.**
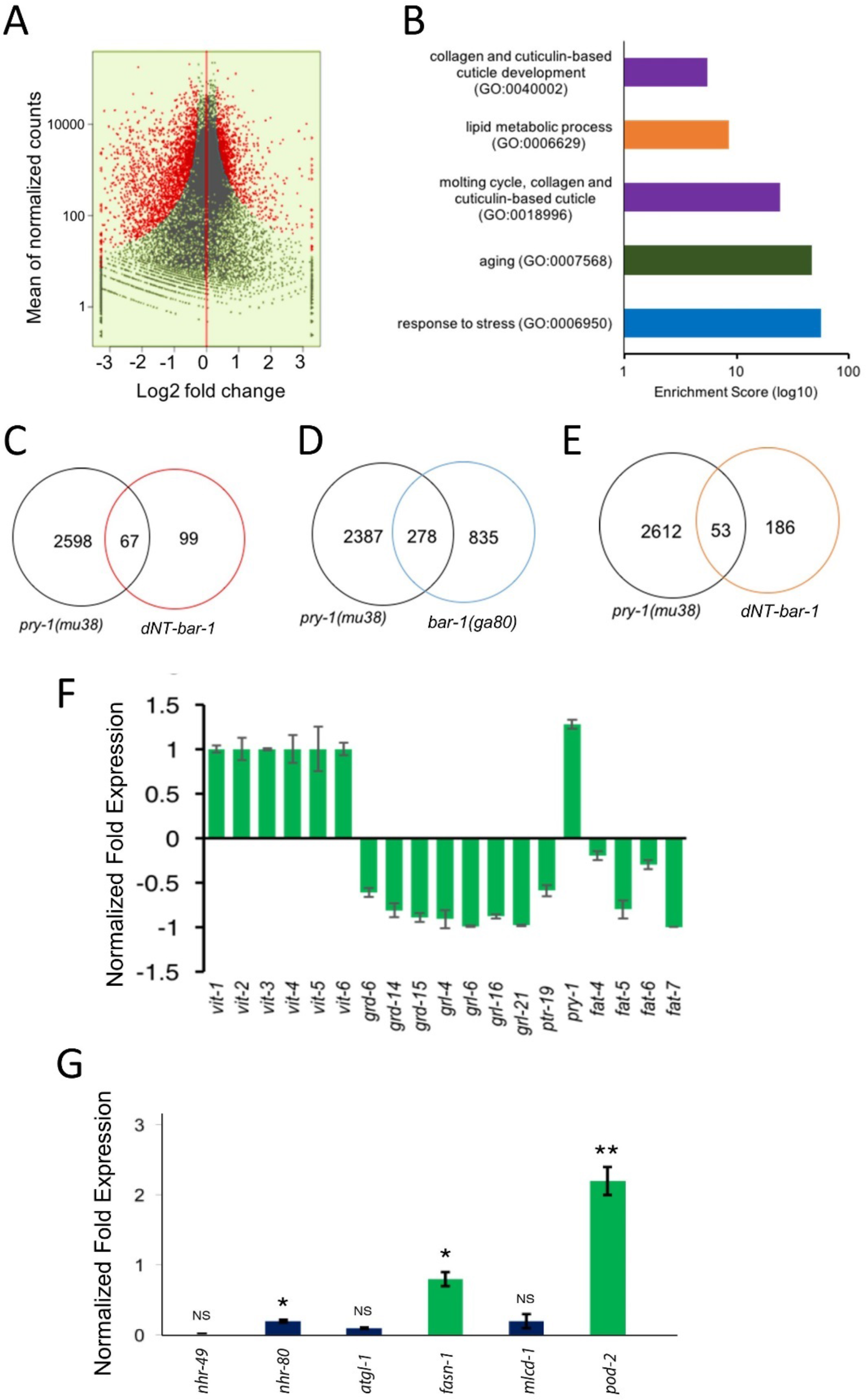
*pry-1(mu38)* transcriptome analysis. **A)** Scatter plots of differentially expressed genes in *pry-1(mu38)*. Red dots mark significantly altered transcripts with a FDR *p*-adj of < 0.05, whereas black dots mark transcripts that are not significantly altered (FDR *p*-values of > 0.05). **B)** Selected GO categories enriched in *pry-1(mu38)* targets are identified by GO Amigo, (*p*-adj < 0.05). (see Table S4 for a detailed list). **C-E)** Venn diagrams showing the overlap between *pry-1(mu38)*, *dNT-bar-1*, and *bar-1(ga80)* transcriptional targets. **C)** Overlap between *pry-1(mu38)* and *dNT-bar-1(28)* (RF:2.5, hyp.geo p < 9.722e-14) **D)** Overlap between *pry-1(mu38)* and *bar-1(ga80)* (27) (RF:1.7, hyp.geo p < 5.325e-19) **E)** Overlap between *pry-1(mu38)* and *dNT-bar-*1(29) (RF:1.5, hyp.geo p < 0.002) **F)** qPCR analysis of mRNA for PRY-1 target genes in the *pry-1(mu38)* mutant. Data represent mean ± SEM of 3 biological repeats. \**p* < 0.001. **G)** Expression of fat metabolism-related genes in *pry-1* mutant. Error bars represent standard error. \**p* < 0.05, \*\**p* < 0.01.

We next carried out gene ontology (GO) analysis (*www.geneontology.org*) to investigate the processes affected in *pry-1(mu38)* animals. Genes with altered expression were found to be enriched in GO terms associated with “determination of adult lifespan”, “aging”, “response to unfolded protein”, “oxidation-reduction process”, “metabolism”, “stress response and cell signaling”, “steroid hormone mediated signaling”, “lipid metabolic processes”, and “cellular response to lipids” (Fig. 1B; a complete list is provided in Table S4). This suggests that *pry-1* plays a role in stress response, lipid metabolism, and lifespan regulation. We also observed enrichment in neuron-related GO terms such as “axon”, “synapse”, “synaptic transmission”, and “neuron development”, which was expected from the requirements of *pry-1* in neuronal development (3). Other categories included “molting cycle”, “regulation of transcription”, “DNA-template”, and the “reproductive process”. In addition to these known categories, the dataset included numerous non-annotated genes (Table S3) whose functions remain uncharacterized.

In particular, aging-related genes and gene families were overrepresented in our dataset (122 in total, 46% upregulated and 54% downregulated) (Table S3). These included genes that encode extracellular matrix proteins such as collagens (51) and cuticulin (14), stress-response factors including glutathione *S*-transferase (23), heat-shock proteins (12), cytochrome P450s (29), insulin signaling-related molecules such as insulin-like peptides (4) and DOD (downstream of *daf-16*) (4), and yolk lipoprotein VIT/vitellogenin (6). Of the 16 collagen genes that are essential for lifespan extension and mediated by dauer signaling (26), 54% were present in our dataset (*col-13*, *141*, *144*, *176*, *180*, *61*, *65*, *89*, *97*) (hyp.geo *p*-value 6.12 e−05).

Notably, several components of Hedgehog (HH) signaling are downregulated in *pry-1(mu38)* animals including warthog genes *wrt-1* and *wrt-9*; groundhog-like genes *grl-1*, *grl-4*, *grl-5*, *grl-6*, *grl-7*, *grl-13*, *grl-16*, and *grl-21*; hedgehog-like genes, *grd-3*, *grd-5*, *grd-12*, *grd-14*, and *grd-15*; and patched-related genes, *daf-6*, *ptr-1*, *ptr-13*, *ptr-16*, *ptr-19*, *ptr-2*, *ptr-20*, *ptr-21*, *ptr-22*, *ptr-23*, and *ptr-8*. This suggests that hedgehog-signaling is affected by *pry-1(mu38)*. Some of these genes were also recovered earlier in *bar-1* transcriptome studies (27–29), discussed further below. The *ptc* and *ptr* genes promote molting and the trafficking of proteins, sterols, and lipids (26, 30). In support of such function, we found molting and other cuticle-related defects in *pry-1* mutants (e.g., rollers, defective alae, and weaker cuticle) (Mallick *et al*., unpublished, manuscript in preparation). These data are consistent with the role of WNT signaling in cuticle development (28).

Moreover, alterations in the expression of some of the Wnt pathway components were noted as well. For example, *pry-1* was upregulated 1.3-fold (Table S3, Fig. S1G). Although such upregulation of *pry-1* has not been reported previously in *pry-1* mutants, prior studies have shown that Axin constitutes a target of WNT signaling and its expression is increased in overactivated WNT backgrounds (5, 28, 29, 31). Together with our finding, this result suggests that the positive regulation of Axin by the canonical WNT signaling pathway represents a conserved mechanism in eukaryotes. Other WNT pathway components that were differentially expressed in *pry-1(mu38)* included *mom-2/wnt* (1.5-fold increase), *cfz-2/fz* (1.7-fold decrease), *lin-17/fz* (1.6-fold increase), and *pop-1/tcf* (1.7-fold increase) (Table S3).

A comparison of *pry-1* targets with three previously reported WNT transcriptome microarray datasets, *bar-1(null)* (27) and constitutively active *bar-1* (*dNT-bar-1*, a small deletion that removes part of the *N*-terminus) (28, 29), revealed shared genes and families. Of the two *dNT-bar-1* studies, one was specific to the vulva and seam cells (32) and the other involved whole animal analysis (28). A total of 12% (29), 18% (27), and 30% (28) *bar-1* targets overlapped with our *pry-1* set (Fig.1C-E, Table S5). The shared targets included genes involved in cuticle synthesis (*col* and *cutl* families), defense response, embryo development, oxidation-reduction processes, and proteolysis. The *hh* family members were also shared between the *pry-1* and *bar-1* targets. Altogether, from the three *bar-1* studies mentioned above, 10 *hh* target genes, namely *grl-5*, *grl-10*, *grl-14*, *grl-15*, *hog-1*, *grd-1*, *grd-2*, *grd-12*, *wrt-4*, and *wrt-6* were reported, of which three (*grl-5*, *grl-14*, and *grd- 12*) are present in our *pry-1* target list. The common targets of *bar-1* and *pry-1* (Fig. 1C-E, Table S5) may therefore act downstream of the canonical WNT signaling pathway

### *pry-1*mutants exhibit altered lipid metabolism

Throughout the lifespan of an animal, lipids are persistently mobilized to supply energy demands for growth, cellular maintenance, tissue repair, and reproduction (33, 34). Changes in lipid levels also affect an organism’s ability to survive in stressful conditions (35, 36). Notably, many genes that are involved in the synthesis, breakdown, and transport of lipids are differentially expressed in *pry-1* mutants (Fig. S2 - lipid metabolism). These include vitellogenins (yolk protein/apolipoprotein-like): *vit-1*–*6*; fatty acid transporters: *lbp-1*, *-2*, *-4*, *-5*, *-7*, and *-8*; lipases: *lips-3*, *-4*, *-7*, *-10*, and *-17*; desaturases: *fat-4-6*; elongases: *elo-3* and *-6*; and fatty acid oxidation: *acdh-1*, −6, *-11*, *-23*, *acs-2*, *-11*, and *-17*, *cpt-1*, *-4*, and *ech-9* (Table S2). The expression of *vit* genes and desaturase was measured by qPCR and all but *fat-4* were successfully validated (Fig. S1G) as levels of the latter were decreased by 20%, unlike the 1.5-fold increase observed by RNA-Seq. We also tested another desaturase, *fat-7*, which functions redundantly with *fat-6* (37) but was not present in our dataset. *fat-7* mRNA levels were below the limit of detection (Fig. S1G); thus, all four fat desaturase genes were downregulated in *pry-1(mu38)* animals.

Enrichment of several lipid metabolism genes in the *pry-1* transcriptome led us to examine lipid accumulation in worms. Staining with Oil Red O revealed that the lipid content was less than half in *pry-1(mu38)* one-day-old young adults compared with controls (Fig. 2A, B). Examination of total fat at each larval stage revealed that *pry-1* mutants had lower somatic lipid stores (25–80%) at all stages except for L2 (Fig. 3A). In addition, the lipid distribution was altered such that the staining was mostly restricted to gonadal tissue (Fig. 2C). These results suggested that *pry-1* plays a role in lipid metabolism. Consistent with this model, we found that *pry-1(mu38)* animals laid fewer fertilized eggs and had poor survival upon starvation-induced L1 diapause (Fig. 2E, F).

**Figure 2.**
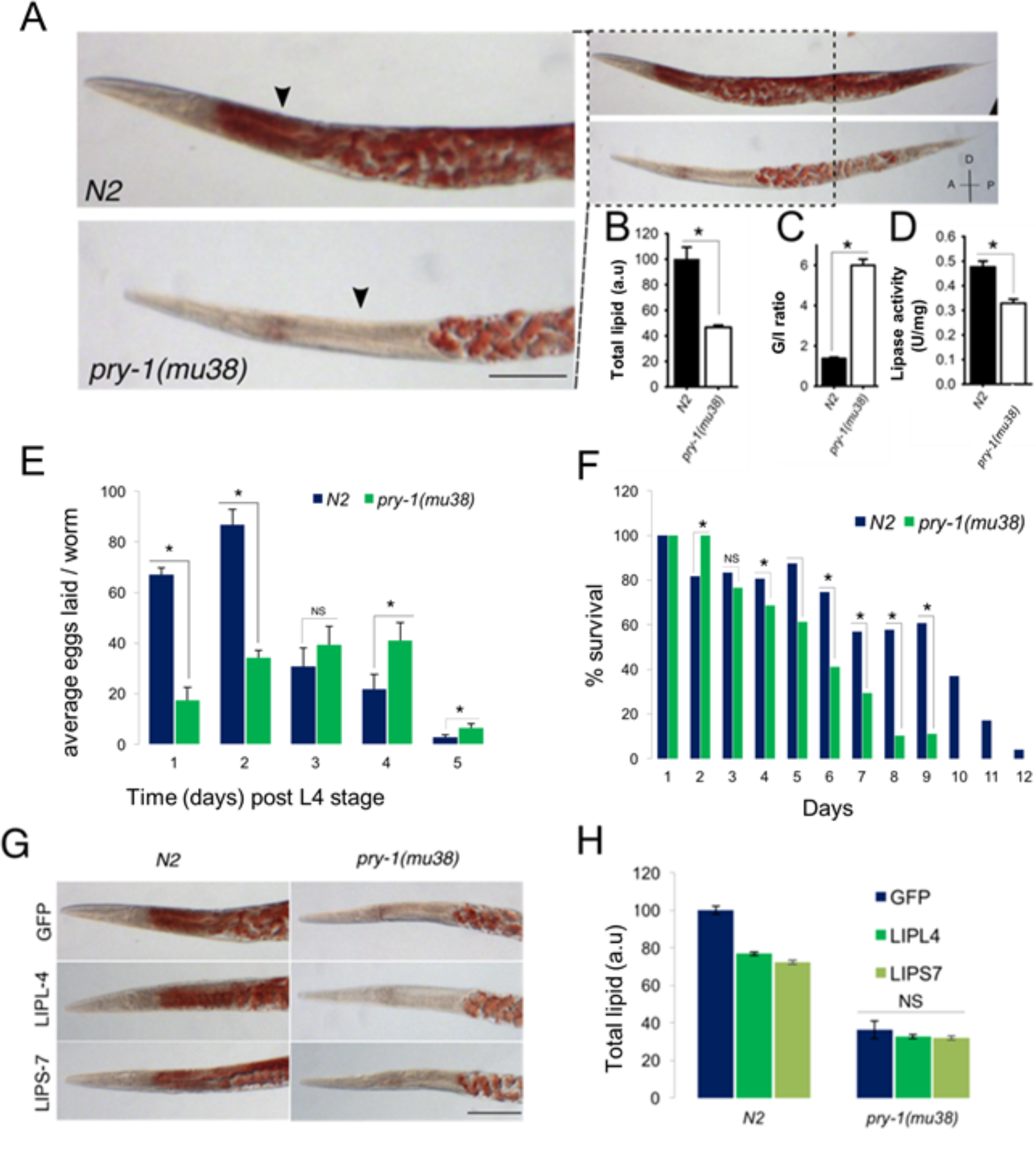
Lipid levels and distribution are altered in *pry-1(mu38)* mutants. Arrowheads indicate intestine and dotted areas gonad. **A)** Representative DIC images of N2 and *pry-1(mu38)*, stained with Oil Red O. D: dorsal; V: ventral; A: anterior; and P: posterior. **B, C)** Quantification of Oil Red O staining. Data are presented as mean; error bars in this and subsequent graphs represent SEM. **C)** Ratio of gonadal to intestine (G/I) lipid. **D)** The lipase activity is decreased in *pry-1(mu38)*. The lipase activity is plotted as activity per mg of protein. **E)** The average number of eggs laid by wild-type and *pry-1(mu38)* animals on different days over the duration of their reproductive period. **F)** *pry-1(mu38)* displayed significant reduction in L1 survival following starvation. Percent survival of L1 larvae in the absence of food has been plotted. Graph represents the average of three independent experiments. **G)** Representative images of N2 and *pry-1(mu38)* following RNAi treatments and Oil Red-O staining. **H**) The histogram shows Oil Red intensity of N2 and *pry-1(mu38)*. Scale bar, 50 µm, \**p* < 0.001 (unpaired t-test).

**Figure 3.**
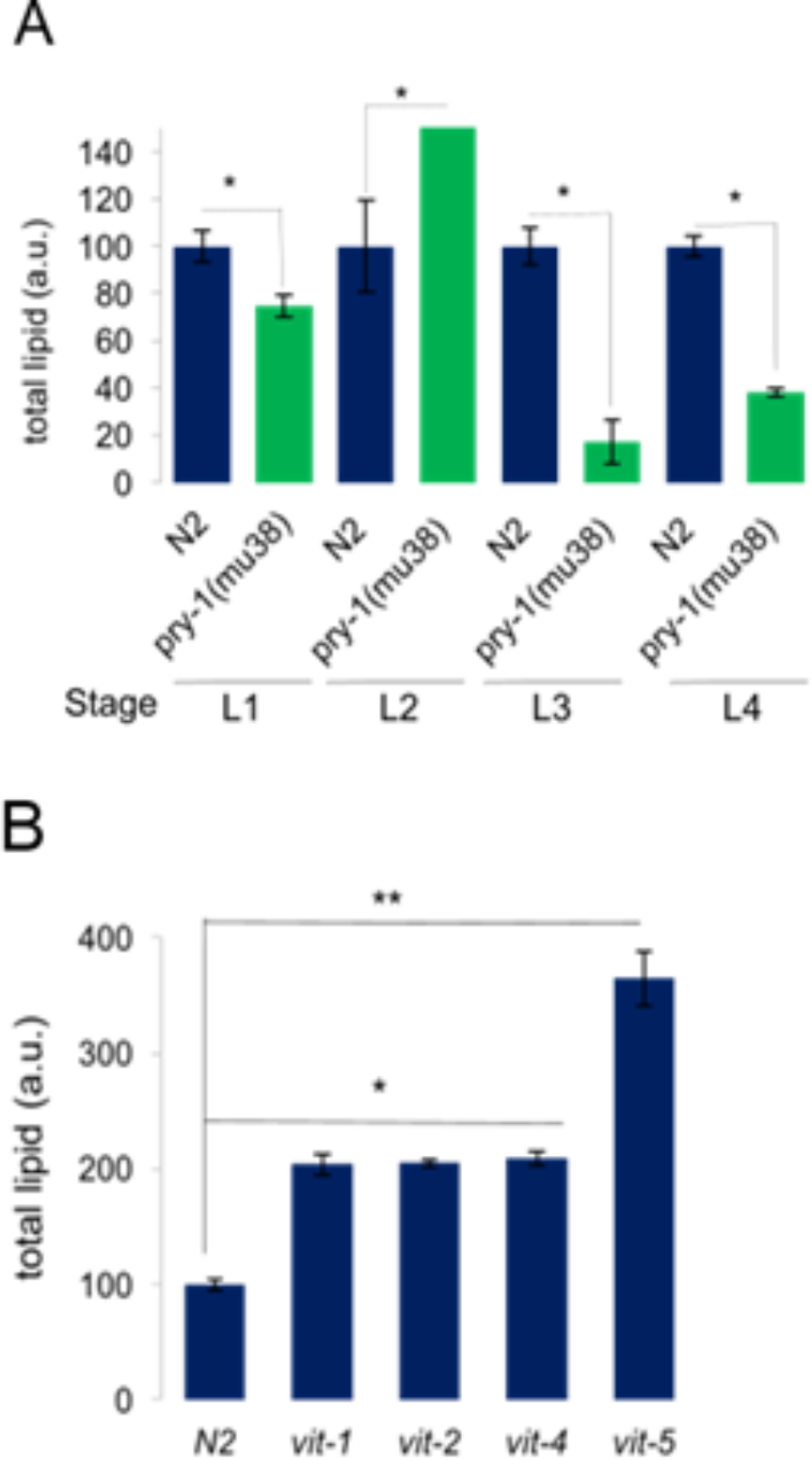
Lipid levels in *pry-1(mu38)* and *vit* mutants. **A)** Quantification of Oil Red O staining intensity in *pry-1(mu38)* and wild-type animals at different developmental stages. Lipids are lower in *pry-1* mutants at all stages except L2. **B)** Quantitation data of Oil Red O staining in N2 and *vit* mutants during the young adult stage. Lipid levels are higher in *vit-1(ok2616), vit-2(ok3211), vit-4(ok2982), and vit-5(ok3239)* animals. Error bars represent mean ± SEM. \**p* < 0.01, \*\**p* < 0.001.

One explanation for the reduced lipid phenotype may be that lipids are being rapidly utilized. However, this is unlikely because several lipases (*lips* family members) were downregulated. We also measured total lipase activity in one-day-old adults from whole worm lysates. As expected, the total lipase activity was 34% lower in the mutant compared with the N2 control (Fig. 2D). Next, we examined lipids in *pry-1(mu38)* animals following knockdown of *lipl-4* or *lips-7*, which comprise lipase genes that regulate the gonad-dependent somatic lipid levels (33, 34, 38) but observed no change in the pattern of lipid distribution (Fig. 2G, H). We concluded that the lower somatic lipids in animals lacking *pry-1* function were not due to increased utilization, raising the possibility of the involvement of other metabolic processes.

### Vitellogenins contribute to lipid metabolism defects in *pry-1* mutants

To understand the molecular basis of low lipid levels in *pry-1(mu38)* worms we focused on the vitellogenin family of genes, whose expression is repressed by *pry-1*. VITs comprise the major yolk proteins in *C. elegans*, which are synthesized in the intestine and mediate lipid transport from the intestine to the gonad during the reproductive period (39). Examination of *vit* levels in *pry-1(mu38)* animals revealed abnormal expression at all developmental stages. Thus, beginning with the L1 stage where all six *vit* genes were upregulated, the number of overexpressed genes was five in L2, two in L3, and zero in L4 stage (Fig. 4).

**Figure 4.**
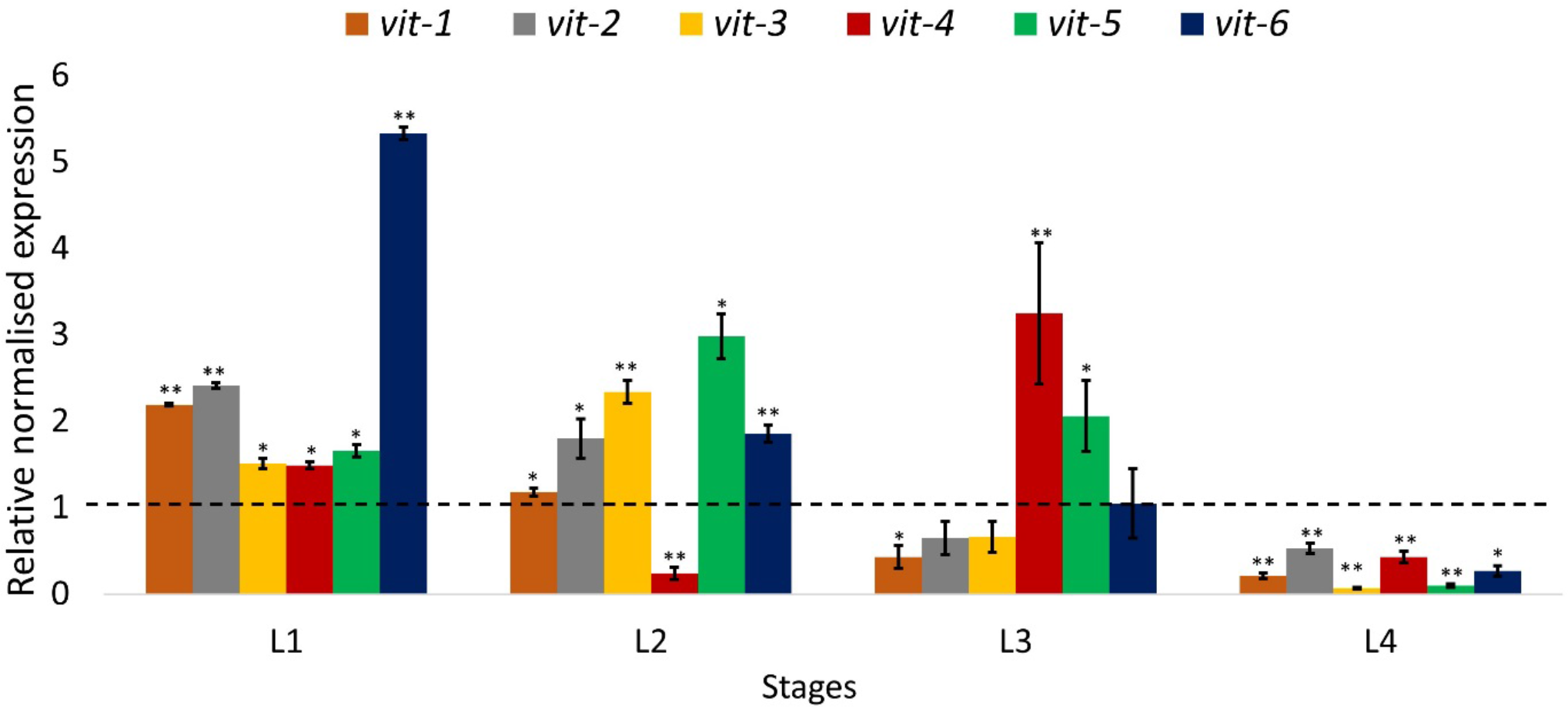
Expression of *vit* genes in *pry-1(mu38)*. **A)** qRT-PCR of *vit* genes at larval stages in *pry-1(mu38)* mutants. Relative normalized expression has been plotted. Error bars represent standard error of the mean (\**p*<0.05 and \*\**p*<0.01 for all mutants compared to wild type).

We next examined lipid contents in worms using mutants and RNAi knockdown of specific *vit* genes. The results revealed altered lipid distributions in all cases such that lipids accumulated at higher levels in somatic tissues (Fig. 3B, Fig. 5A, B). A quantification of overall lipids following *vit* knockdowns revealed a significantly higher accumulation in mutants compared to wild-type (2.5–3 fold and 1.3–1.4 fold, respectively, using GFP RNAi knockdown as a control in each case) (Fig. 5A, B). Lipid accumulation following *vit* knockdown in wild-type has also been previously reported (40). Sequence comparisons indicated that *vit-1* RNAi may also target *vit-2* owing to significant identity in the genomic region used to perform KDs (Table S6), which was validated by qPCR (Fig S3). Similarly, any one of the *vit-3*, *4*, or *5* RNAi could simultaneously effect knockdown of the other two (Table S6). Thus, multiple VITs appear to play roles in the regulation of both the level as well as the distribution of lipids, and their misregulation contributes to lipid metabolism defects in *pry-1* mutants.

**Figure 5.**
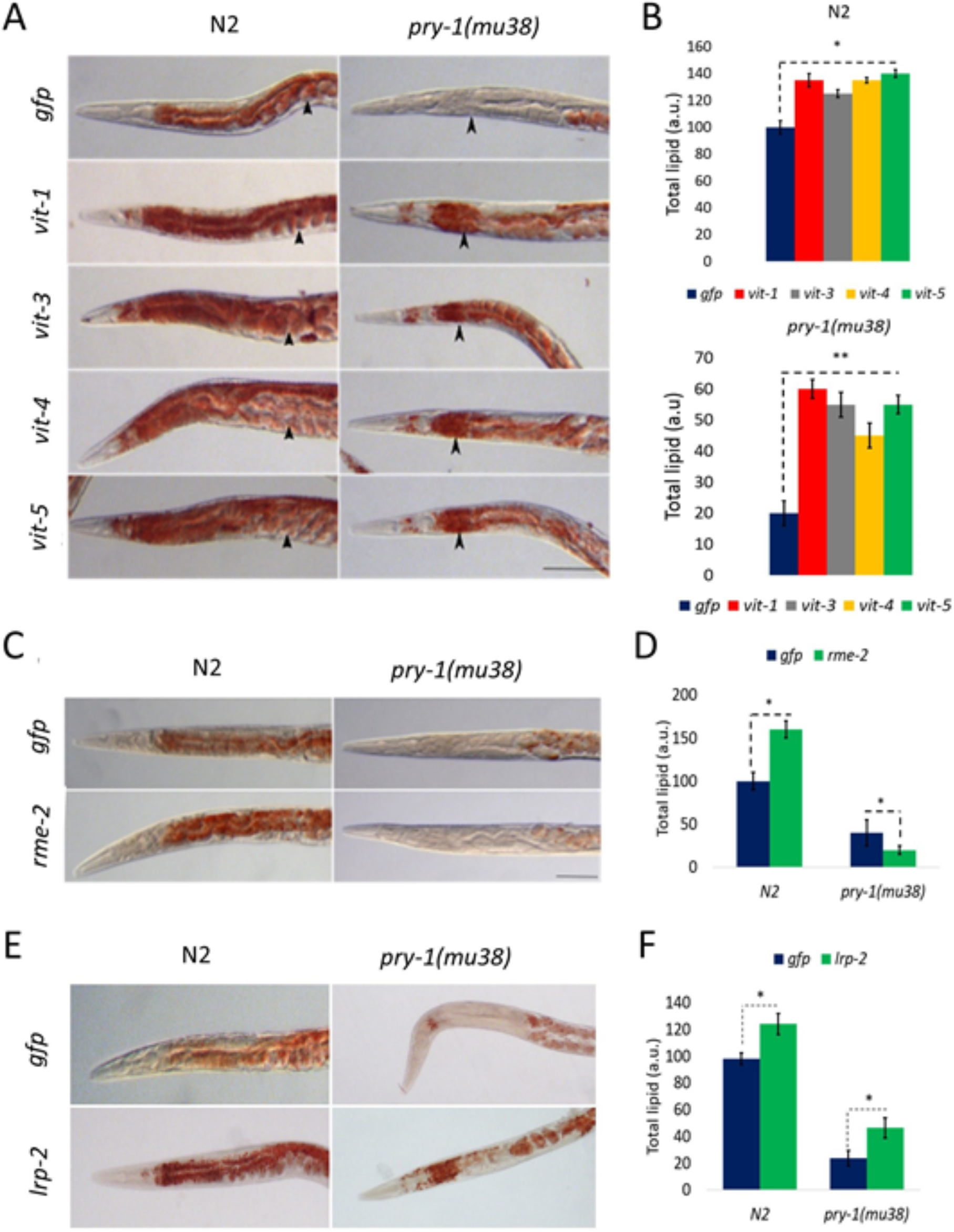
Vitellogenin mediated lipid metabolism in *pry-1* mutants. **A)** DIC micrographs of representative N2 and *pry-1(mu38)* adults stained with Oil Red-O treated with *vit* RNAi. **B)** The corresponding Oil Red O quantifications following each treatment. **C)** Representative images of N2 and *pry-1(mu38)* following *rme-2* RNAi KD. **D)** The histogram shows Oil red intensity of N2 and *pry-1(mu38)*. **E)** Representative images of N2 and *pry-1(mu38)* following *lrp-2* RNAi KD. **F)** The histogram shows Oil red intensity of N2 and *pry-1(mu38)*. (At least 50 animals were quantified in each batch. Two batches done).

### Lipoprotein receptors RME-2 and LRP-2 may not contribute to lipid defects in *pry-1*mutants

To understand how *pry-1* and *vit* genes might function to regulate lipid levels, we examined the involvement of *rme-2* in the *pry-1*-mediated pathway to regulate lipid accumulation as VITs are transported via the RME-2 receptor (39). The knockdown of *rme-2* by RNAi led to intestinal accumulation and ectopic deposition of lipids (Fig. 5C, D) consequent to blockage of yolk protein transport to the developing oocytes (39). Specifically, *rme-2*(RNAi) animals showed an approximately 45% increase in total lipid content such that the gonad-to-somatic ratio was roughly 30% lower compared with that in controls. However, this phenotype was not observed in *pry-1(mu38)* owing to a reduction in lipid levels both in somatic and gonadal tissues (Fig. 5C, D). These results allow us to suggest that VITs act independently of the RME-2 transport mechanism to regulate lipid metabolism in response to *pry-1* signaling. Moreover, as lipid levels are further reduced in *pry-1(mu38)* animals following *rme-2* knockdown, *rme-2* may have an unknown non-vitellogenin-mediated role in lipid accumulation, although other possibilities also exist for such a phenotype.

We also examined the possibility of other VIT-interacting factors that might be involved in PRY-1-mediated regulation of lipid levels. Our transcriptome dataset contained one LDL-like receptor gene, *lrp-2*, which was overexpressed in *pry-1* mutants (Table S3). It was shown previously that *lrp-2* positively regulates yolk protein synthesis (11). To test whether lipid levels are affected by *lrp-2*, RNAi knockdown experiments were performed, which showed a small but significant rescue of the lipid phenotype in *pry-1(mu38)* animals (Fig. 5E, F). However, as *lrp-2* knockdown in wild-type animals also caused a similar increase in lipids, it is unclear whether PRY-1-mediated signaling incorporates LRP-2 function to affect lipid levels.

### Fatty acid levels are reduced in *pry-1* mutants

Among the *pry-1* transcriptome genes predicted to participate in fatty acid desaturation and elongation (5), binding/transport (6), and β-oxidation (14) pathway (2) (Fig. S2), the four Δ9-desaturase enzymes, which show downregulated gene expression in *pry-1* mutants, are required to produce C16:1 and C18:1 monounsaturated fatty acids (Fig. S2). Whereas *fat-5* converts palmitic acid (C16:0) to palmitoleic acid (C16:1n7), *fat-6* and *fat-7* are involved in stearic acid (C18:0) to OA (C18:1n9) conversion (41). Although mutations in single Δ9-desaturase genes cause reduced fatty acid synthesis, they do not give rise to a significant visible phenotype because of a compensatory mechanism; however, double and triple mutants exhibit severe defects, including lethality (41).

Because the expression of all four fat genes depends on the nuclear hormone receptors *nhr-49* and *nhr-80* (25, 42), we determined levels of these NHR transcripts in *pry-1* mutant animals. Although RNA-Seq transcriptome data showed no change in either gene, qPCR revealed that *nhr-80* transcription showed a subtle but significant upregulation whereas *nhr-49* was unchanged (Fig. 1G). Thus, transcriptional regulation of these two NHRs is unlikely to be a mechanism affecting *pry-1*-mediated expression of Δ9-desaturase genes, although we cannot rule out the possibility that activities of one or both may be regulated post-transcriptionally in response to PRY-1 function.

The changes in the expression of Δ9-desaturases may lead to reduced fatty acid synthesis in *pry-1(mu38)* animals. To investigate this, we quantified lipid levels using a gas chromatography-mass spectrometry (GC-MS) approach. The results showed that whereas the relative ratios of fatty acids in *pry-*1 mutants were normal, the absolute level of each species was significantly reduced (Fig. 6A, Fig. S4). The result agrees with the overall low lipid levels in *pry-1(mu38)* animals and together supports the important role of PRY-1 signaling in lipid metabolism in *C. elegans*.

**Figure 6.**
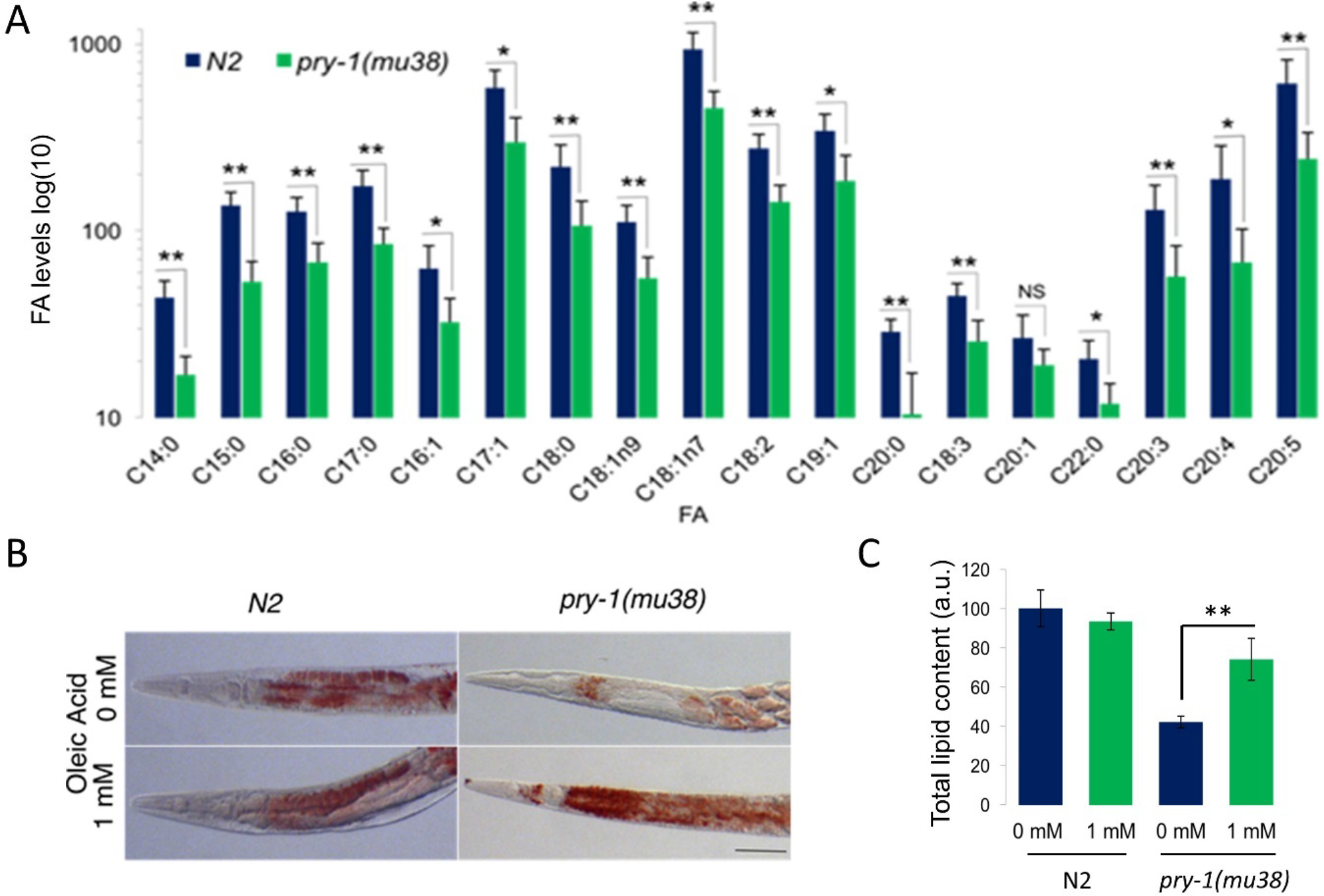
GC-MS analysis of fatty acids in *pry-1* mutants and partial rescue of lipid defects following OA treatments. **A)** Total FA levels of selected fatty acid species expressed in log_10_ value as determined by GC-MS analysis. The *pry-1* mutants have low levels of FA. Error bars represent the standard deviation. Significant differences between wild type and *pry-1* mutant are marked with stars, * *p*< 0.03, ** *p* < 0.015. **B)** Lipid staining of N2 and *pry-1(mu38)* adults supplemented with 1 mM OA. **C)** Quantification of total lipid, *p* < 0.001.

### OA (18C:1n9) supplementation partially rescues the somatic depletion of lipids in *pry-1* mutants

OA constitutes one of the fatty acid species that showed 50% reduced levels in our GC-MS analysis. OA is required for fatty acid metabolism and is synthesized by *C. elegans* as it cannot be obtained through the normal *E. coli* (OP50) diet. OA acts as a precursor for the synthesis of polyunsaturated fatty acids and triacylglycerides, which are used for fat storage (24). The addition of exogenous OA as a fat source has been shown to rescue several fat-deficient mutants including *fat-6* and *fat-7* by restoring their fat storage, resulting in improved fertility and increased locomotion (37). Moreover, the addition of OA in *sbp-1*, *fat-6*, and *fat-7* animals fully rescued defects in satiety quiescence (41–43). We therefore reasoned that supplementation of OA may improve lipid levels in *pry-1(mu38)* mutants. Treatment with 1 mM OA resulted in the restoration of lipids in animals lacking *pry-1* function (up to 2-fold higher compared with the untreated control, Fig. 6B, C). No significant changes were seen in the gonadal lipid levels, suggesting that lipid metabolism in the gonad was unaffected.

## DISCUSSION

### *pry-1* regulates expression of genes involved in lipid metabolism

To understand the mechanism of *pry-1/Axin* function, in the present study we performed transcriptome profiling on a *pry* mutant. The analysis revealed altered expression of many genes including those that affect hypodermis, stress-response, aging, and lipid metabolism. The hypodermal-related genes included collagens, cuticulins, and hedgehogs. Previously, expression of some of the hedgehog genes was found to be altered in *bar-1* mutants (27, 29). Considering that cuticular defects are observed in *bar-1* (28, 29) and *pry-1* mutants (Mallick *et al*., manuscript in preparation), and that hedgehog family members play roles in cuticle shedding and the formation of alae (30), these results led us to suggest that a genetic pathway involving *pry-1* and *bar-1* may interact with hedgehogs for normal cuticle development.

One of the key findings of our *pry-1(mu38)* transcriptome analysis was the enrichment of genes related to lipid metabolism. In particular, we found that multiple lipogenic and lipolytic genes exhibited altered expression. For example, all four fatty acid desaturases (∆5 and ∆9 desaturases) were downregulated in *pry-1* mutants. Whereas single *fat* gene mutants affect fatty acid composition without altering overall lipid levels, double mutants have a low lipid level (37, 41), suggesting that *pry-1* positively regulates fatty acid synthesis. With regard to lipolytic genes, such as those involved in beta-oxidation, changes in gene expression between peroxisomal and mitochondrial beta-oxidation genes showed an opposite trend (4 of 5 upregulated and 8 of 11 downregulated, respectively). This may indicate selective utilization of long-chain over short-chain fatty acids by the *pry-1* pathway. We also observed that all four lipases (*lips* family) were downregulated including *lips-7*, which was previously shown to be involved in lifespan extension and the maintenance of lipid levels (38). Although *lips-7* did not alter the *pry-1(mu38)* phenotype, it remains to be seen whether *pry-1* regulates any, or all, of the remaining three *lips* gene(s) to modulate lipids.

In addition to lipogenic and lipolytic genes, several lipid transporters are also present in the *pry-1(mu38)* transcriptome, including two lipid-binding proteins (*lbp-5* and *lbp-8*; both downregulated), six lipoproteins (*vit-1* to *-6*; all upregulated), and a LDL-like receptor protein (*lrp-2*). Knockdown of *lbp-5* and VITs negatively affect lipid storage (44), which further emphasizes the important role of *pry-1* in the maintenance of lipids and suggests that *pry-1*-mediated signaling is involved in the utilization of lipids for energetics as well as signaling mechanisms.

### PRY-1-mediated lipid metabolism involves vitellogenins

Reduced lipids may affect tissue function and physiology in different ways; for example, owing to altered membrane structure and compartmentalization, altered signaling, reduced energy demands, and impact on autophagy. The Oil Red O staining of *pry-1(mu38)* showed a severe reduction in lipid content with a marked decline in the somatic lipid storage. It is worth mentioning that a similar phenotype was also observed in *dNT-bar-1* animals that carry a constitutively active form of BAR-1 (Fig. S5). Together these findings suggest that the misregulation of Wnt signaling affects lipid metabolism in *C. elegans*.

One possibility for reduced lipids in *pry-1* and *bar-1* mutants may be due to changes in the enzymatic activities of lipases and lipid desaturases (24, 41, 42). However, *pry-1* mutants showed no increase in total lipase activity. Additionally, knockdowns of *lipl-4* (lysosomal lipase) and *lips-7* (cytosolic lipase), both of which negatively regulate lipid levels (33, 34), in *pry-1(mu38)* had no observable effect. Thus, selective and rapid lipid catabolism does not appear to be a factor in lipid depletion in the absence of *pry-1* function.

We then investigated the role of VIT proteins, which are the distant homologues of human apolipoprotein B (45), in maintaining lipid levels. As major yolk proteins, VITs are involved in somatic mobilization of lipids to the developing germline. Moreover, previous studies demonstrated that reducing VITs in wild-type animals increases both lifespan and lipid accumulation, with overexpression having an opposite effect in long-lived mutants (40, 46). *pry-1* mutants showed misregulation of all six *vit* genes such that they were overexpressed in L1 but declined thereafter with all being downregulated by the L4 stage. As expected, the low lipid phenotype of *pry-1* mutant animals was suppressed by knocking down *vit* genes (*vit-1/2* and vit*-3*/*4*/*5*), providing evidence that VITs play an important role in PRY-1-mediated lipid metabolism. We have also shown that such a role of VITs may not utilize lipoprotein receptors RME-2 (VIT transporter) or LRP-2 (VIT synthesis). Overall, these findings along with the role of VITs in regulating lipid levels (40), allow us to propose that PRY-1-mediated signaling involves VITs to regulate processes that depend on energy metabolism and lipid signaling (33, 34).

Although it remains unclear how VITs participate in *pry-1* signaling, one possibility may involve downregulation of the autophagy pathway. Manipulating VIT levels has been shown to affect the lifespan by altering autophagy (40). Autophagy is a complex process that involves multiple enzymes to recycle cellular contents by converting them into usable metabolites. Despite the *pry-1* transcriptome not containing known autophagy-related candidates, the pathway may still be involved. This could be tested by examining the roles of specific autophagosome genes in lipoprotein synthesis and autophagy, which in turn should reveal a link with *pry-1*-mediated lipid metabolism.

Another possibility for low lipid levels in animals lacking *pry-1* function may be due to reduced fatty acid synthesis. In agreement with this, the expression of three conserved stearoyl-CoA desaturases, *fat-5*, *fat-6*, and *fat-7*, which are involved in the synthesis of monounsaturated fatty acids such as oleic acid (OA) (24, 41, 42) was reduced in *pry-1* mutants. Moreover, our GC-MS analysis of fatty acid composition revealed that *pry-1* is needed to maintain normal levels of every fatty acid species analyzed. A global reduction in fatty acids owing to the loss of *pry-1* function may affect processes that require utilization of lipids such as aging. However, the relationship between lipid levels and lifespan is not well understood. It is likely that rather than absolute levels, the quality of lipids may be more important for cellular processes (36). We investigated this possibility using OA, one of the species involved in fatty acid signaling. Exogenous treatment with OA restored lipid levels in *pry-1* mutants. Thus, *pry-1* may play a role in maintaining the levels of beneficial fatty acids. This then raises the question of how the *pry-1*-*vit*-mediated pathway might affect lipid levels. It is possible that lipid synthesis and storage processes are compromised, which is supported in part by the reduced expression of desaturases; however, additional mechanisms are also likely to be involved, such as reduced conversion of acetyl-CoA to saturated fatty acid (palmitate), lower synthesis of diglycerides, and increased peroxisomal beta oxidation (Fig. S2). It would therefore be worthwhile to examine these possibilities in future studies.

Our study provides the first evidence of PRY-1/Axin function in lipid metabolism. The involvement of lipids in age-related disorders in humans, as well as animal models, is well documented. Genetic and acquired lipid diseases are associated with loss of subcutaneous fat, accumulation of visceral and ectopic fat, and metabolic syndromes such as insulin resistance, glucose intolerance, dyslipidemia, and hypertension (47). In addition, Yang et al. showed that Axin expression in mice contributes to an age-related increase in adiposity in thymic stromal cells (48). Although our data show that PRY-1/Axin is not likely to affect fat storage, whether Axin family members play roles in any of these lipid-related diseases remains unknown. Therefore, the findings that PRY-1/Axin is necessary for the regulation of lipid levels provide a unique opportunity to investigate the role of Axin signaling in age-related lipid metabolism.

## ACKNOWLEDGMENTS

We are grateful to Brian Golding for providing server space, the Paul Sternberg lab for help with RNA-Seq computational analysis, Hendrik Korswagen for the *pry-1p*::PRY-1::GFP plasmid, and the Don Moerman lab for *pry-1* CRISPR alleles. Some of the strains were obtained from CGC, which is funded by the NIH Office of Research Infrastructure Programs (P40OD010440). We thank Lesley Macneil and anonymous reviewers for comments on previous versions of the manuscript, Jessica Knox for assistance with initial quantitative real time-PCR experiments, and Gupta lab members for helpful discussions. This work was supported by an NSERC Discovery grant to BG.

## AUTHOR CONTRIBUTION

BG conceived and supervised the entire project. AR, AM and BG designed experiments and wrote the manuscript. AR and AM performed experiments. The authors declare no conflict of interest.

## SUPPLEMENTAY DATA

**Figure S1.**
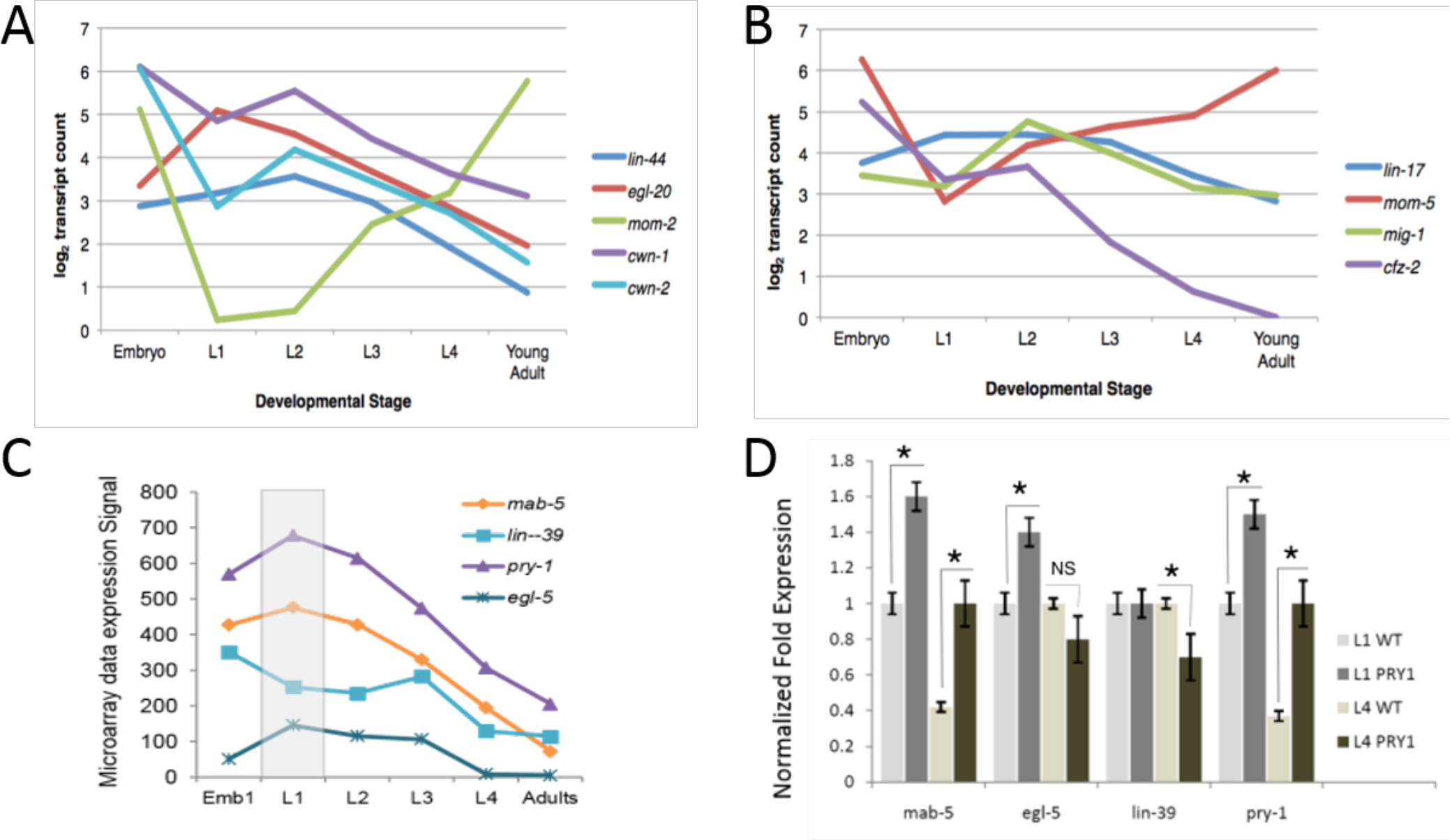
Expression profile of Wnt ligands, receptors, and target genes. **A-C)** Developmental expression patterns of known Wnt ligands, receptors and target genes from published microarray sources (see Methods). **D)** qPCR validations of selected Wnt target genes during the L1 and L4 stages.

**Figure S2.**
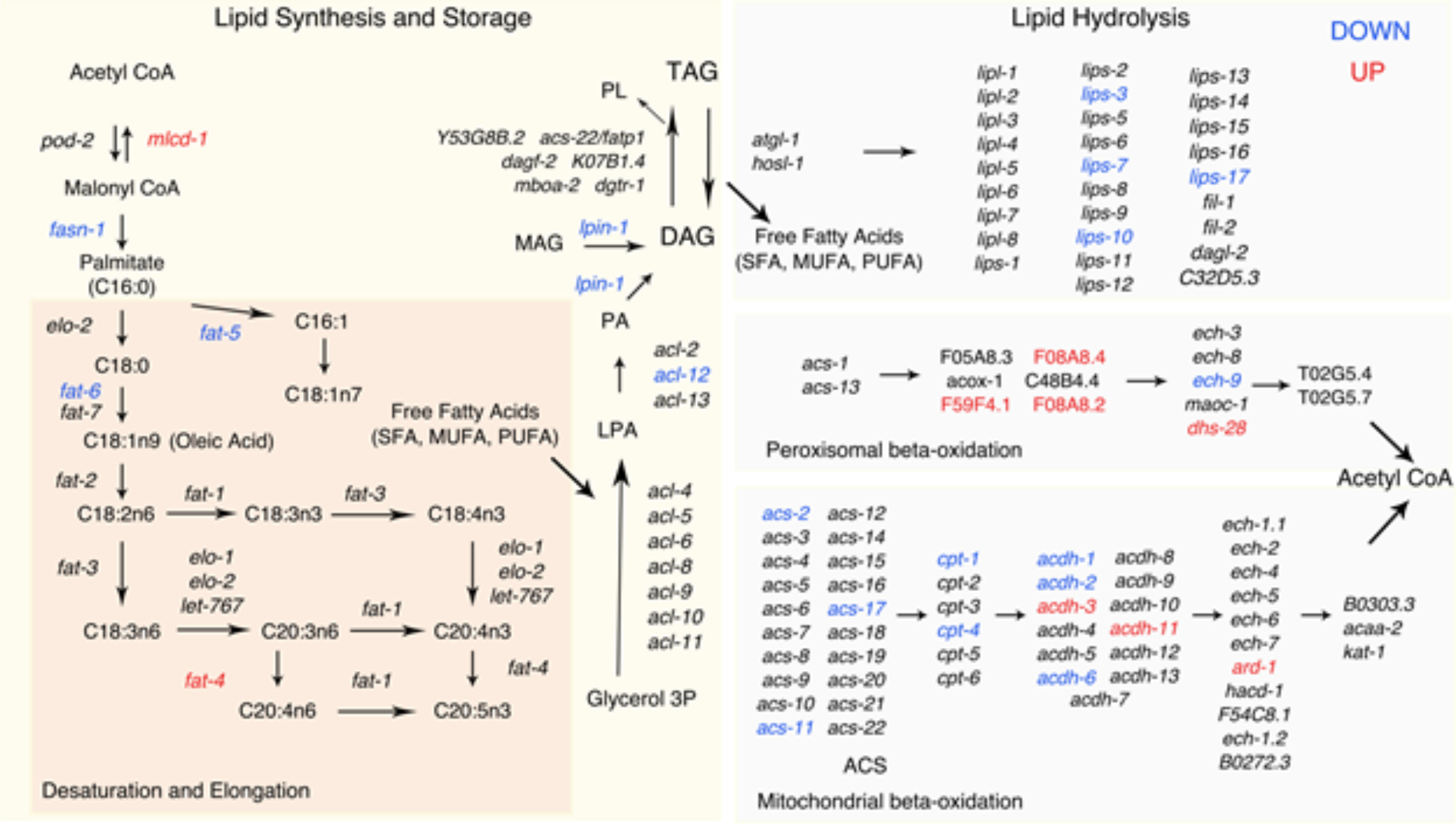
Overview of lipid metabolism and genes with altered expression in *pry-1(mu38)*. The lipid anabolic and catabolic pathway is adapted from a previously published study (Amrit *et al*. 2016a; Amrit *et al*. 2016b). Lipid anabolic processes involve initiation, desaturation and elongation of fatty-acid (FA), followed by triglyceride (TAG) formation. Initiation involves conversion of Acetyl CoA to the saturated fatty-acid (SFA) Palmitate (C16:0). Elongase (*elo*) and desaturase (*fat*) enzymes act on Palmitate to modify it to long chain mono- and poly-unsaturated fatty acids (MUFAs and PUFAs, respectively). MUFAs and PUFAs are collectively termed as free fatty acids (FFAs). The FFAs are linked with glycerol 3-phosphate (Glycerol 3P) to produce lysophosphatidic acid (LPA) and phosphatidic acid (PA). PA and monoglycerides (MAG) serve as building blocks of diglycerides (DAG) synthesis. DAGs are converted into neutral lipids (TAGs). Lipid catabolism begins with the breakdown of TAGs into DAGs by ATGL-1, and other lipases and lipase-like enzymes (abbreviated as ‘*lipl’* and ‘*lips’*) to release FFAs. FFAs are further broken down to Acetyl CoA through peroxisomal- and mitochondrial-β-oxidation and release energy. Putative genes with enzymatic activity that are involved in lipid metabolism are shown at the appropriate step. Genes with altered expression in *pry-1(mu38)* are highlighted in blue (DOWN) and red (UP).

**Figure S3.**
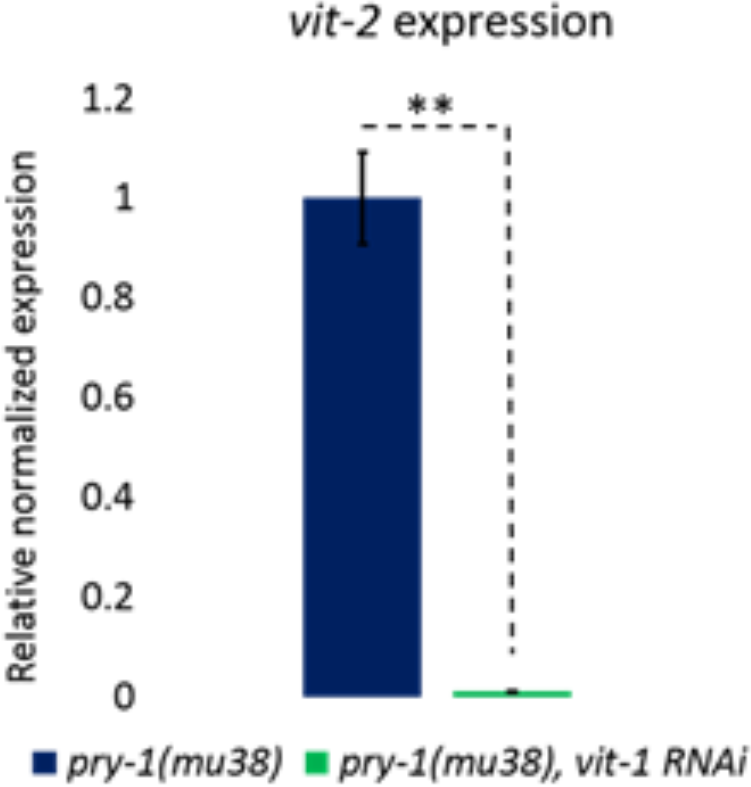
*vit-1* RNAi Knocks down *vit-2* transcript level. **A)** qPCR analysis of *vit-2* in the *pry-1(mu38)* day 3 mutants after adult specific *vit-1* RNAi KD. Data represent mean ± SEM of 3 biological repeats. \**p* < 0.01.

**Figure S4.**
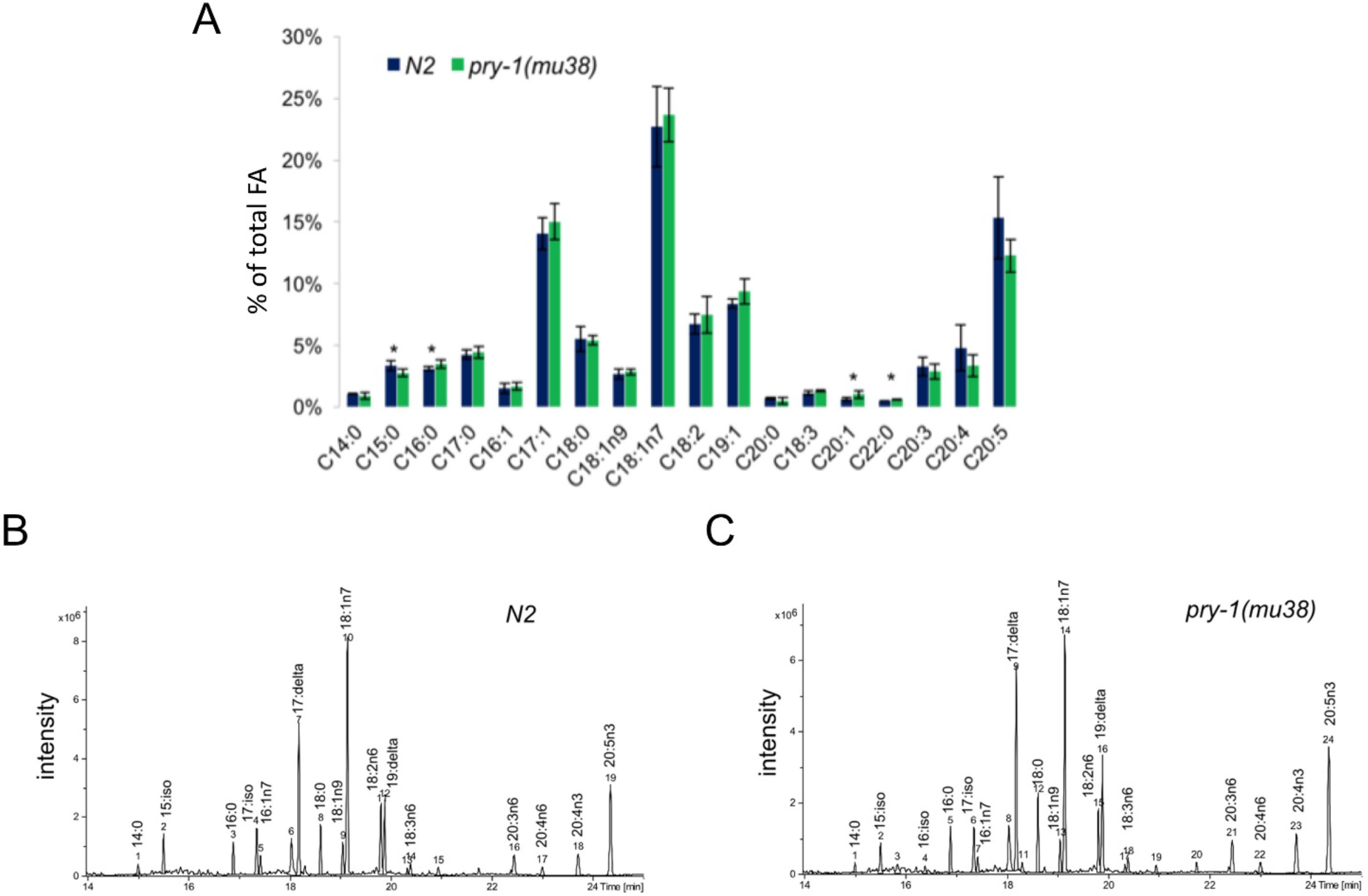
Relative fatty acid abundance in *pry-1(mu38)*. **A)** Relative abundance of selected fatty acid species expressed in percentage of total fatty acid as determined by GC-MS analysis. *pry-1* mutants have marginally lower levels of C15:0, C16:0 and higher levels of C20:1, C22:0 than N2 (marked with star, *p* < 0.05). Error bars represent the standard deviation. **B, C)** A representative GC-MS Total Ion Chromatogram (TIC) traces of populations of the N2 and *pry-1(mu38)* worms, respectively. The peaks corresponding to fatty acid species are identified.

**Figure S5.**
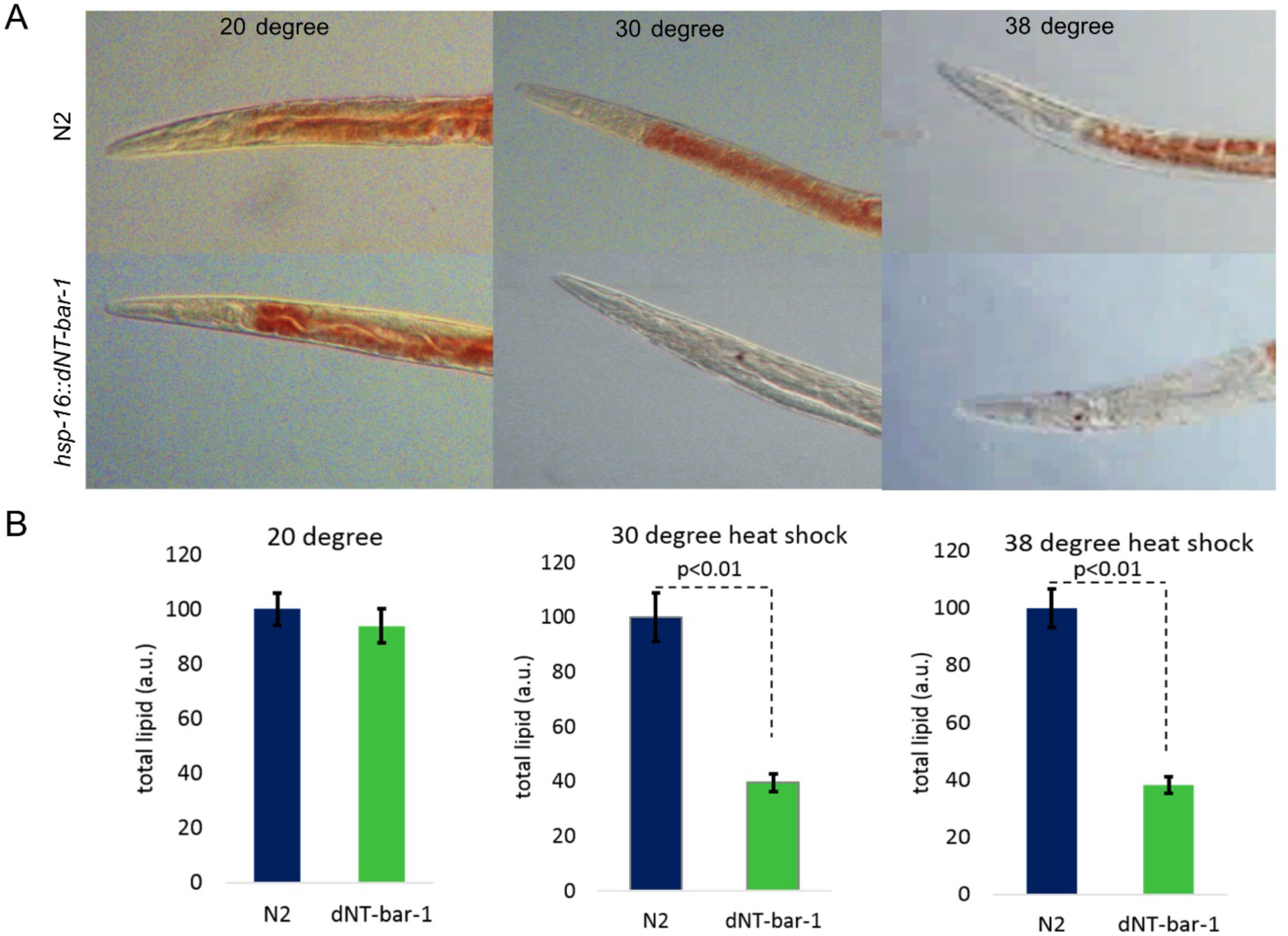
Lipid staining and quantification in *dNT-bar-1* animals. **A)** Representative DIC images of N2 and *dNT-bar-1* at 20 °C, after heat shock at 30 °C for 12hrs and 38 ^0^C for 30 minutes, stained with Oil Red O. **B)** Quantification of total lipids in dNT-bar-1 animals after heat shock treatments. (2 trials, n>50 for each trial; *p<0.01* for all mutants compared to wild type).

**Table S1.**
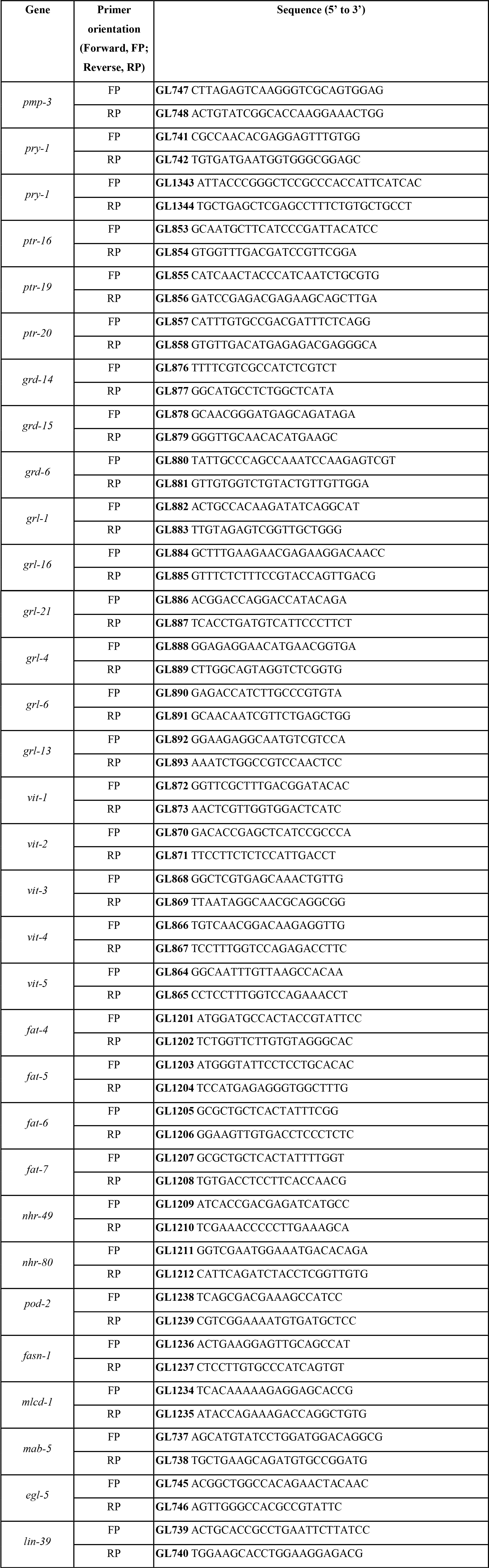
List primers used in this study.

**Table S2.**
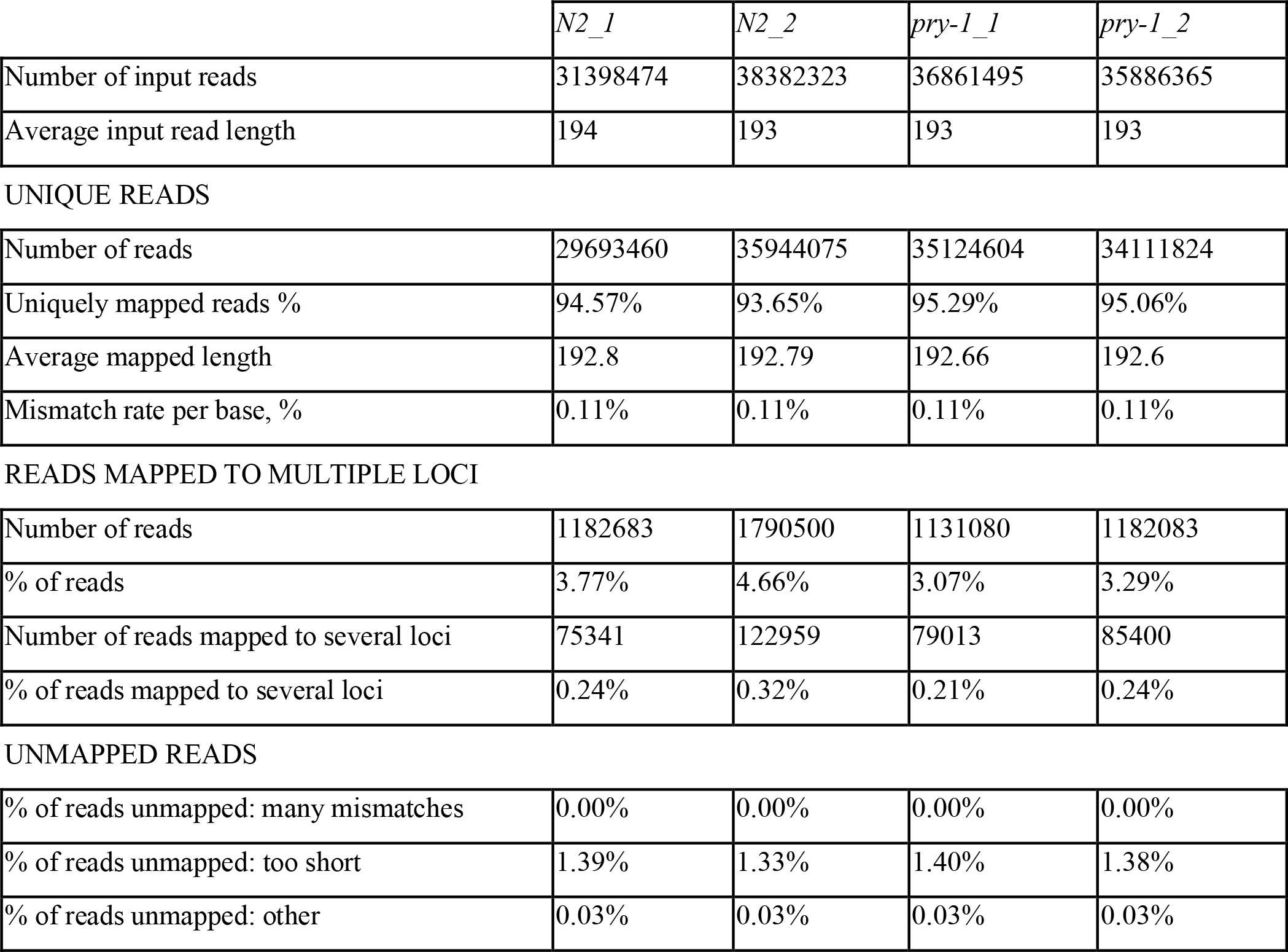
The number mRNA transcripts mapped to the *C. elegans* genome.

|  | *N2_1* | *N2_2* | *pry-1_1* | *pry-1_2* |
| --- | --- | --- | --- | --- |
| Number of input reads | 31398474 | 38382323 | 36861495 | 35886365 |
| Average input read length | 194 | 193 | 193 | 193 |
| UNIQUE READS |  |  |  |  |
| Number of reads | 29693460 | 35944075 | 35124604 | 34111824 |
| Uniquely mapped reads % | 94.57% | 93.65% | 95.29% | 95.06% |
| Average mapped length | 192.8 | 192.79 | 192.66 | 192.6 |
| Mismatch rate per base, % | 0.11% | 0.11% | 0.11% | 0.11% |
| READS MAPPED TO MULTIPLE LOCI |  |  |  |  |
| Number of reads | 1182683 | 1790500 | 1131080 | 1182083 |
| % of reads | 3.77% | 4.66% | 3.07% | 3.29% |
| Number of reads mapped to several loci | 75341 | 122959 | 79013 | 85400 |
| % of reads mapped to several loci | 0.24% | 0.32% | 0.21% | 0.24% |
| % of reads unmapped: many mismatches | 0.00% | 0.00% | 0.00% | 0.00% |
| % of reads unmapped: too short | 1.39% | 1.33% | 1.40% | 1.38% |
| % of reads unmapped: other | 0.03% | 0.03% | 0.03% | 0.03% |

**Table S3.**An Excel spreadsheet listing differentially regulated genes.

**Table S4.** An Excel spreadsheet showing GO-term enrichment.

**Table S5.** An Excel spreadsheet showing transcriptome comparison.

**Table S6.** Conservation of *vit* gene sequences used in RNAi experiments.

| gene targets | RNAi |  |  |  |
| --- | --- | --- | --- | --- |
|  | *vit-1* | *vit-3* | *vit-4* | *vit-5* |
| *vit-1* | 100% | ns | 80% (98bp) | 80% (123bp) |
| *vit-2* | 94% (1025 bp) | ns | 80% (99bp) | 81% (123bp) |
| *vit-3* | ns | 100% | 99% (2421bp) | 94% (1125bp) |
| *vit-4* | ns | 98% (2364bp) | 100% | 97% (1075bp) |
| *vit-5* | ns | 97% (2267bp) | 98% (2397bp) | 100% |
| *vit-6* | ns | ns | ns | ns |

ns: no significant identity observed.

